# Spared Nav1.8-Positive Nociceptors Drive Persistent Tactile Hypersensitivity After Sciatic Nerve Crush Injury in Mice

**DOI:** 10.1101/2025.11.09.687306

**Authors:** Sang Wook Shim, Yoon Kyung Lee, Dahee Roh, Kihwan Lee, Hyoung Woo Kim, Seog Bae Oh

## Abstract

Peripheral nerve injury can lead to chronic mechanical hypersensitivity, yet the severity and persistence of pain are strongly influenced by the extent of axonal damage. Notably, partial sciatic nerve crush injury (PCI) produces persistent tactile hypersensitivity despite a less severe anatomical insult than full crush injury (FCI), yet the identity and post-injury state of the fibers that persist after PCI remain unclear. To define sensory neuron populations contributing to PCI-induced tactile hypersensitivity, we combined fiber-specific transgenic labeling (Thy1-YFP for Aβ mechanoreceptors and Nav1.8-tdTomato for nociceptors) with pharmacological silencing using QX-314 coapplied with TRPV1 (capsaicin) and TLR5 (flagellin) agonists to selectively manipulate fiber subtypes. At day 7 after PCI, Nav1.8^+^ nociceptive terminals were still detectable in the hind paw. On day 30, acute silencing of TRPV1^+^ afferents transiently reduced mechanical hypersensitivity, indicating nociceptor activity in its maintenance. Whole-cell patch-clamp recordings of retrogradely labeled DRG neurons showed that remaining medium-diameter neurons exhibited reduced rheobase and increased action potential firings in response to step current injections. Besides, electrical stimulation of nociceptive fibers increased pERK expression in the spinal dorsal horn, indicating enhanced nociceptive signaling after PCI. Early ablation of TRPV1^+^ fibers with high-dose capsaicin during degeneration phase prevented the subsequent development of long-term tactile hypersensitivity. Collectively, our results suggest that spared nociceptors after PCI remain sensitized even during nerve repair, driving long-term tactile hypersensitivity. Targeting these spared nociceptive fibers after nerve injury may offer a potential strategy for preventing chronic pain associated with traumatic nerve injury.

**Graphical Summary:** 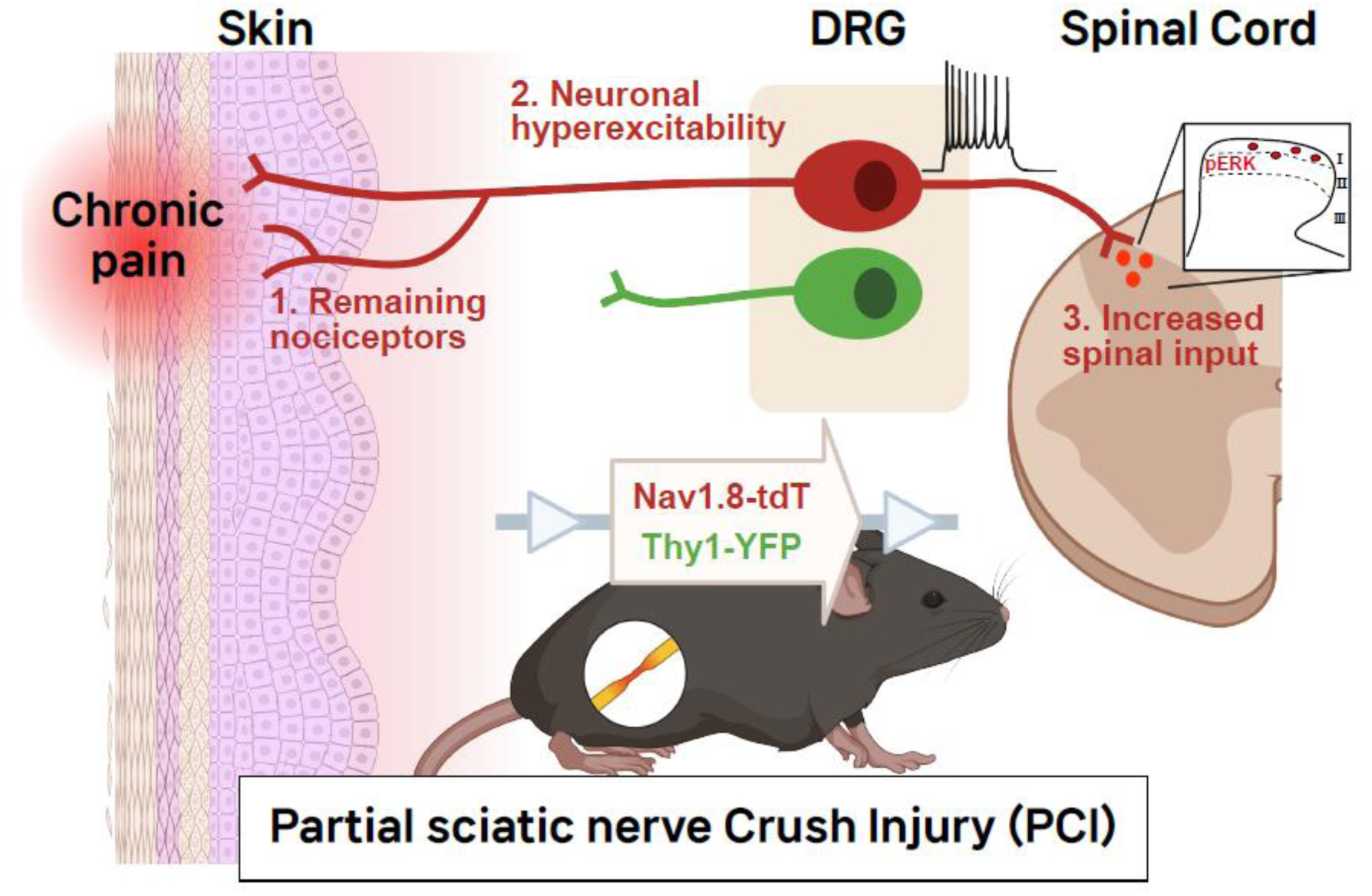

## 1. Introduction

Neuropathic pain after peripheral nerve injury can persist for months to years and imposes substantial personal and societal burdens [20; 54]. Nerve injuries often result in a complex mixture of damaged and intact neurons [25; 51]. Both injured and uninjured primary afferent neurons can contribute to pain [14; 31; 83; 84], and the molecular and functional heterogeneity of dorsal root ganglia (DRG) neurons complicates pinpointing which neuronal populations drive hypersensitivity [50; 75]. This heterogeneity yields fiber-class–specific responses to injury: small-diameter nociceptors (Aδ/C) and large myelinated Aβ mechanoreceptors differ in vulnerability and post-injury plasticity across lesion types and severities [21; 56]. Accordingly, identifying which afferent classes are spared or injured, and how their intrinsic excitability and target innervation are remodeled, remains crucial for linking specific injury contexts to pain development.

We recently reported that partial sciatic nerve crush injury (PCI) results in delayed, yet persistent tactile hypersensitivity compared with full crush injury (FCI), despite a less severe anatomical lesion [42]. This paradoxical outcome was associated with the selective sparing of small-diameter sensory fibers, suggesting that these surviving neurons may drive long-term pain. However, the specific identity of these spared neuronal populations and whether they undergo functional changes that could promote chronic hypersensitivity remain unclear.

Accumulating evidence from various nerve injury models indicates that spared sensory fibers contribute to persistent pain through both functional and structural mechanisms. Functionally, these fibers often exhibit ectopic spontaneous firing and increased excitability in DRG neurons [14; 83], thereby promoting central sensitization by strengthening synaptic transmission in the dorsal horn of the spinal cord [5]. Structurally, histopathological studies reveal aberrant sprouting and mistargeting of spared nociceptors into inappropriate peripheral targets, further amplifying nociceptive signaling [29; 31].

Therefore, we aimed to elucidate the role of spared fibers in driving long-term tactile hypersensitivity following PCI. To address this, we combined fiber-specific transgenic labeling (Thy1-YFP and Nav1.8-tdTomato) with behavioral pharmacological silencing (QX-314 co-applied with TLR5 or TRPV1 agonists) approaches. We examined innervation patterns of the sensory fibers and measured hypersensitivity in the hind paw skin. We also assessed intrinsic excitability of retrogradely labeled DRG neurons by whole-cell patch-clamp recordings, and quantified spinal activation by pERK quantification after fiber-biased electrical stimulation. Our findings show that, unlike FCI, PCI preserves a population of hyperexcitable Nav1.8^+^ nociceptors, and that eliminating these fibers prevents development of tactile hypersensitivity. These findings highlight an important role of spared nociceptors in PCI-induced tactile hypersensitivity and advance understanding of nerve crush-associated pain mechanisms.

## 2. Materials and Methods

### 2.1 Animals

All procedures were approved by the Institutional Animal Care and Use Committee (IACUC) at Seoul National University (protocol code: SNU-220627-3-1, SNU-220111-3-2). C57BL/6J mice (Jackson Laboratory, JAX, stock 000664) were purchased from DooYeol Biotech (Seoul, Republic of Korea). Thy1-YFP-16 mice (RRID: IMSR_JAX:003709) [28] were a kind gift from Dr. Pilhan Kim (KAIST). Nav1.8-cre mice (RRID: IMSR_JAX:036564) [58], Rosa26LSL-tDTomato/+ mice (RRID: IMSR JAX:007909), Trpv1tm1Jul/J (RRID: IMSR_JAX:003770) [12] were purchased from Jackson Laboratory. All experiments were performed on adult mice of both sexes aged 6 to 10 weeks. Mice were bred in a specific pathogen free facility and transferred to a conventional room at least one week before experiments. Animals were maintained on 12-hour light/dark cycle, housed 5-6 mice per cage on wood chip bedding and provided with standard laboratory feed and water ad libitum. Animal husbandry was performed in accordance with the Guideline for Animal Experiments by the Korean Academy of Medical Sciences, the National Institutes of Health Guidelines for the Care and Use of Laboratory Animals.

### 2.2 Surgery

Mice were placed under isoflurane anaesthesia (2% induction) in 100% oxygen (300 mL/min; SomnoSuite; Kent Scientific, CT, USA). Mice were maintained on isoflurane anaesthesia using a face mask (40 mL/min; 2% isoflurane) and body temperature was maintained with a warm pad. Both partial and full crush injury were performed with sample protocol in previous research. In brief, the right thigh was shaved and sanitized with iodine. Incision was made mid-thing length and the sciatic nerve exposed. The nerve was carefully freed from connective tissue and crushed for 15 s using ultra-fine hemostat (Cat no.13020-12, Fine Science Tools, CA, USA). In case of PCI, 30 μm spacers using aluminum foil (Seong Won Cooking Foil, Cheonan, Republic of Korea) were applied on the tip of the ultra-fine hemostat and crushed for 15 s. Full crush was performed in the same manner without spacer. For the sham, the sciatic nerve was exposed without crush injury. The wound was closed with 6-0 silicon-coated silk and the skin incision was closed with 9-mm skin clips (Mikron Precision Inc., Gardena, CA, USA). Animals were placed under an infrared radiator until the recovery from anesthesia without additional analgesia.

### 2.3 Behavioral testing

#### 2.3.1 Evoked pain behaviors

The animals were placed singly in transparent acrylic cylinder on an elevated mash floor. Mice were habituated 3 hours for 3 consecutive days and an hour prior to the following test. Sensory function was measured using pinprick test. Briefly, the response to pinprick stimuli was scored out of 10, as previously described [47]. Mediolateral side hind paw was divided into 5 regions, and an Austerlitz insect pin (size 000, Fine Science Tools, CA, USA) was applied twice to each region. The contralateral hind paw was tested as a positive response. To assess tactile sensitivity, 50% withdrawal threshold was measured using up-down method with a series of *von Frey* filament (0.02-2 g, Stoelting Wood Dale, IL, USA) applied between the central footpads and lateral plantar surface of each paw. The 0.4-g filament was used as the first stimulus.

Thermal sensitivity was measured using a Hargreaves radiant heat apparatus (IITC life Science, CA, USA). Mice were placed in transparent acrylic container (8.5 x 8.5 x 17-cm) above the transparent glass floor, with the temperature maintained at 30 °C. The basal paw withdrawal latency was set between 10 to 15 seconds (auto intensity 25, cut-off time 20 sec). Both hind paws were stimulated for 4 times with intervals of 15-20 minutes. The average latency was calculated based on four individual stimulations.

#### 2.3.2 Selective silencing of sensory fibers using QX-314

We used a cocktail of QX-314 and capsaicin/or flagellin to selectively block Aβ fiber and nociceptive fibers, as previously described [8; 9; 85]. To block cutaneous Aβ fibers, single intra-plantar injection of either vehicle (20 μL of 6 mM QX-314 in sterile saline) or ultrapure flagellin (0.3 μg in 20 μL of vehicle; InvivoGen, CA, USA) was applied into the ipsilateral hind paw. To block cutaneous TRPV1-expressing nociceptor, single intra-plantar injection of either vehicle (20 μL of QX-314 in sterile saline with 10% EtOH) or capsaicin (10 μg in 20 μL of vehicle; 360376; Sigma, MO, USA) was applied into the ipsilateral hind paw. All injections were administrated using an insulin syringe under blind condition. The von Frey test was performed prior to the injection, as well as at 1- and 24-hour post-injection.

#### 2.3.3 Capsaicin ablation of nociceptive nerve terminals

We used capsaicin (360376; Sigma, MO, USA) to selectively ablate nociceptive nerve terminals, as previously described [4; 77]. A single focal injection of either vehicle (20 μL, 10% EtOH in sterile saline) or capsaicin (10 μg in 20 μL of vehicle) subcutaneously injected into the ipsilateral hind paw at day 3 following PCI. The same dose of capsaicin or vehicle was applied on ipsilateral hind paw on day 15 after the crush injury when pain was already generated.

### 2.4 Electrophysiology

#### 2.4.1 Retrograde tracer labeling

Under brief isoflurane anesthesia, 6 µL of 0.5% DiI (Cat no. D282, Molecular Probes, OR, USA) in DMSO was injected into the plantar surface of ipsilateral hind paw at 6 days post-injury [42]. The tracer was injected in the middle of the lateral side of the hind paw within the sciatic nerve innervating area. And 30 days after injury, the lumbar L3-L5 dorsal root ganglia (DRGs) were dissected out for patch clamp recordings.

#### 2.4.2 Primary DRG neuron culture

Primary culture of DRG neurons was performed, as previously described [23]. We used the ipsilateral L3-L5 DRGs from each group of n=3-6 mice. DRG were acutely dissected on ice-cold Hank’s balanced salt solution (HBSS) (with 20 mM HEPES) and digested for 1 hour in collagenase A (1 mg/mL, Roche, Risch-Rotkreuz, Switzerland) and dispase II (2.4 U/mL, Roche, Risch-Rotkreuz, Switzerland) at 37 °C. Then, 0.25% trypsin (Gibco; Thermo Fisher Scientific, MA, USA) in HBSS was applied for 7 minutes. DRG were mechanically dissociated by trituration with fire-polished Pasteur pipette in Dulbecco’s Modified Eagle Medium (DMEM) containing DNase I (125 U/mL, Sigma-Aldrich, MO, USA). The cell suspension was then carefully layered over on 15% bovine serum albumin in Ham’s F-12 (Welgene Inc., Gyeongsan, Republic of Korea) to isolate neurons. Cells were resuspended with Neurobasal medium (Gibco; Thermo Fisher Scientific, MA, USA) containing B-27 supplement (Gibco; Thermo Fisher Scientific, MA, USA), L-glutamine (Sigma-Aldrich, MO, USA), and penicillin-streptomycin (Gibco; Thermo Fisher Scientific, MA, USA). Isolated DRG neurons were plated on a 10-mm diameter coverslip coated with 100 μg/mL poly-D-lysine (PDL, Sigma-Aldrich, MO, USA).

#### 2.4.3 Whole-cell patch clamp recordings

Whole-cell patch clamp recording of DRG neurons was performed, as previously described [18]. Coverslips containing adherent DRG cells were put in the recording chamber and attached to the stage of microscope (Olympus BX51WI, Tokyo, Japan), and DiI-labeled neurons were identified by their fluorescence. Neuronal activities were recorded with a Multiclamp 700B amplifier (Axon Instruments; Molecular Devices, CA, USA) and Digidata-1550B (Axon Instruments, CA, USA). The patch electrode was pulled from BF150-86-10 borosilicate glass capillaries (Sutter Instruments, CA, USA) with P-700 microprocessor-controlled puller (Sutter Instruments, CA, USA) and contained in intracellular solution (containing [in mM] 123 K-gluconate, 18 KCl, 10 NaCl, 3 MgCl_2_, 2 ATP-Na_2_, 0.3 GTP-Na, and 0.2 EGTA, adjusted to pH 7.3 with KOH). Resistance of glass micropipettes was kept at 4 to 6 MΩ. A bath solution (2 mM Ca^2+^-HBSS containing [in mM] 140 NaCl, 5 KCl, 1 MgCl_2_, 2 CaCl_2_, 10 HEPES, and 10 glucose) was adjusted to pH 7.4 with NaOH.

### 2.5 Electrical stimulation for fiber type specific activation

Sham-operated and nerve-injured mice were deeply anesthetized with sodium pentobarbital before the electrical stimulation (100 mg/kg, *i.p.*). Electrodes were securely positioned on the right hind paw’s plantar surface using tape. A series of transcutaneous nerve stimuli were delivered at 2000 Hz, 250 Hz, and 5 Hz to the right hind paw for one minute, targeting Aβ-, Aδ-, and C-fibers, respectively [45; 52; 53]. The respective current intensities applied were 1000 μA for the 2000 Hz stimulus and 2000 μA for the other frequencies. Following a two-minute interval after stimulation, the L4 spinal cord was harvested for the assessment of phosphorylated extracellular signal-regulated protein kinase (pERK) levels.

### 2.6 Single cell RT-PCR

Single cell RT-PCR was performed, as previously described [48; 49]. Before collecting, neurons were examined for DiI signals (red) under a fluorescence microscope. During the cell collection procedure, we continuously flowed a freshly prepared 2 mM Ca^2+^/Na^+^ solution (bath solution). The medium-diameter DiI-labeled neurons were manually collected using a glass micropipette filled with a pipette solution containing RNase inhibitor (RNaseOUT, Invitrogen; Thermo Fisher Scientific, MA, USA). We also collected the bath solution without cells as the negative control for each batch. Complementary DNA (cDNA) of collected neurons was synthesized by reverse transcriptase (Superscript III, Invitrogen; Thermo Fisher Scientific, MA, USA) and used for nested PCR amplifications using nested primer pairs. All primers were designed as exon junction primers to avoid residual genomic DNA contamination (listed in **Supplemental Table 1**).

### 2.7 Immunohistochemistry

#### 2.7.1 Sample preparation

Mice were anesthetized by intraperitoneal injection pf sodium pentobarbital (100 mg/kg) and perfused with 0.1M PBS followed by 4% PFA in 0.1M PBS buffer. Lumbar spinal cord, L4 DRG were dissected from the spinal column and postfixed with the same fixative at 4 °C overnight. Tissues were cryoprotected in 30% sucrose for more than 5 days at 4 °C and embedded in O.C.T. compound (Tissue-Tek, SAKURA, CA, USA). Skin in mediolateral hind paw was isolated using a 3-mm biopsy punch and post-fixed 15 minutes at RT followed by 30% sucrose solution for cryoprotection at 4 °C. Samples were cryostat sectioned at 10 μm for DRG, 12 μm for hind paw skin, 30 μm for spinal cord with a cryostat microtome (Leica, etzlar, Germany) and mounted onto Superfrost glass slides. All sectioned samples were stored at -80 °C deep freezer. The sections were blocked and permeabilized with 5% host serum and 0.3% PBST (Triton-X 100) for 1 hour at room temperature. Tissues were incubated overnight at 4 °C with primary antibodies. After the rinsing of the sections with PBS, secondary antibodies and DAPI (1 mg/mL, Cat no. D9542, Sigma, MO, USA) were applied to the sections for 1 to 2 hours at room temperature. Antibody dilution was prepared using 1% host serum in 0.03% PBST (Triton-X 100) (listed in Supplemental Table 2), and PBS was used for washing the antibodies.

#### 2.7.2 Whole tissue clearing and immunostaining

For the whole mount staining, hind paw skin was dissected out after PBS perfusion followed by 4% PFA in 0.1 M PBS buffer. The tissues were post-fixed in 4% PFA at 4 °C overnight and stored in PBS. Fat and connective tissues were thoroughly removed to facilitate antibody penetration. Tissues were blocked and permeabilized with blocking solution (20% DMSO, 5% normal host serum in 1% Triton X-100 in PBS; PBSt) overnight at 4 °C and then primary antibody were diluted in blocking solution for 3 days at 4 °C, as previously described [40]. Tissues were washed twice, each for an hour, with PBS followed by 1% PBSt and then incubated with secondary antibodies diluted in blocking solution for 3 days at 4 °C. Tissues were washed twice, each for an hour, with PBS followed by 1% PBSt. Finally, tissues were cleared in OPTIMUS solution overnight at 4 °C and mounted onto confocal dish with OPTIMUS solution [44]. All incubations were done on a shaker.

### 2.8 Confocal imaging and analysis

Images were acquired using a confocal microscope (Zeiss LSM980, Oberkochen, Germany). For imaging whole mounted hind paw skin, 23-25 z-stack sections on 10 ⅹ lens with 6 tile scans (interval 6 μm; 120-150 μm in total depth) and exported as maximum intensity projection. Low-magnification whole-mounted imaging was used to evaluate overall denervation and re-innervation patterns covering entire plantar region. For 20 ⅹ images, 7-8 z-stack sections (interval 6 μm; 36-42 μm in total depth) were acquired in three random regions per tissue samples to observe specific innervation to Merkel cells by each type of fiber (Thy1^+^ *vs.* Nav1.8^+^). The number of Merkel and percent of co-labeling with Thy1+ and Nav1.8+ fibers were manually counted within the skin region targeted in each experiment. And for the automatic analysis, we used IMARIS software to calculate shortest distance and overlapped volume of Thy1+ and Nav1.8+ fiber with K8+ Merkel cells. Total branch lengths of individual nerve terminals were automatically measured using Image J software (NIH).

### 2.9 Statistics

For *in vivo* experiments, biological unit of interest is the number of mice. For patch clamp experiments, the biological unit of interest is the cells (i.e. number of DRG neurons). All statistical analyses were performed using IBM SPSS Version 26.0. The normal distribution of the data was analyzed with a Shapiro-Wilk test before further applying parametric or non-parametric statistical analyses. Unpaired Student’s *t*-test and one-way ANOVA were used for normally distributed data. When data did not follow a normal distribution, the Mann-Whitney *U* test was used to compare the data between groups, and the Friedman test was used to formally detect differences between groups across repeated measures. Data were also tested for homogeneity of variance using Levene’s test. Investigators performing the behavioral test, quantitative histological staining and morphometric analyses were blinded either to the surgery or treatment group. Treatments were assigned to littermates at random by an independent observer. Data represents as mean ± standard error of the mean (SEM). \**p* < 0.05 was considered significant.

## 3. Results

### 3.1 Selective silencing of TRPV1-positive nociceptors relieves PCI-induced hypersensitivity

We first sought to identify the specific sensory fiber populations responsible for PCI-induced hypersensitivity. We used QX-314 (N-ethyl-lidocaine), a membrane impermeable sodium channel blocker [8; 9; 85]. To selectively silence fiber classes, QX-314 was co-applied with either flagellin (a TLR5 agonist biasing A-fiber entry) or capsaicin (a TRPV1 agonist biasing C-fiber entry) (**Supplemental Fig. 1**).

Consistent with our prior works [42; 43], tactile hypersensitivity developed by 15 days after PCI and persisted through day 30 (**Fig. 1A**). Next day (day 31 post-PCI), we co-administered QX-314 with each fiber-specific agonist (**Fig. 1B**). Co-application of QX-314 with capsaicin transiently relieved tactile hypersensitivity an hour post-injection, compared to QX-314 alone (**Fig. 1C**), whereas co-administration of QX-314 with flagellin had no effect on pain behavior (**Fig. 1D**). To assess the direct role of TRPV1 channel, we used Trpv1-knockout (KO) mice [12]. The tactile hypersensitivity developed similarly in both wild-type and KO mice, indicating that the TRPV1 channel itself is not required for the development of pain-like behavior (**Fig. 1E**). Collectively, these findings suggest a contribution of TRPV1^+^ nociceptor populations to PCI-induced hypersensitivity, while indicating that TRPV1 channel activity itself is not necessary for the development.

**Figure 1.**
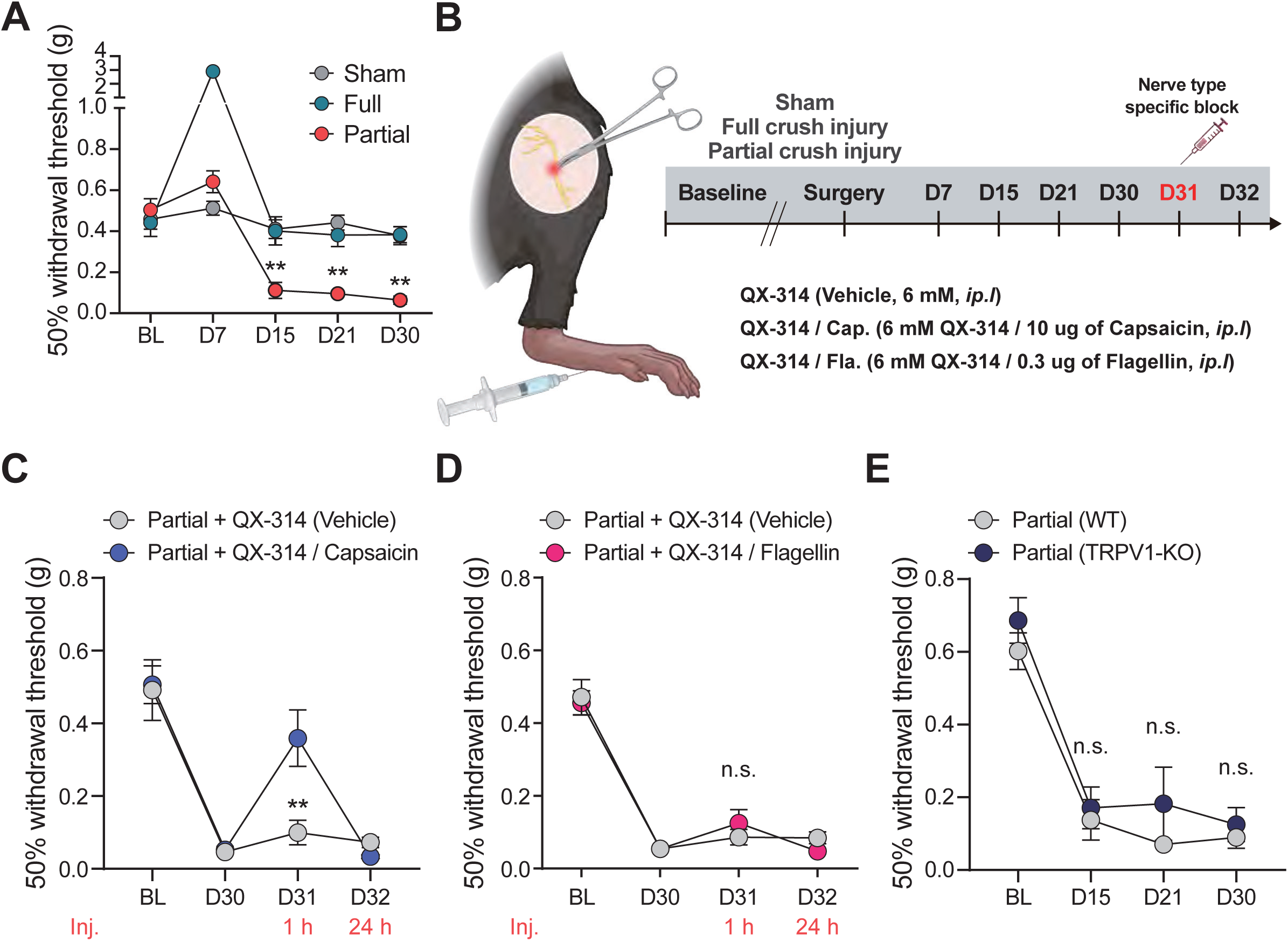
TRPV1+ fiber blockade by co-application of capsaicin/QX-314 inhibits mechanical hypersensitivity following partial crush injury. **(A)** 50% withdrawal threshold of ipsilateral paw following partial, full crush and sham surgery group. Data represent mean ± SEM (n=5-8 mice per group, Sham *vs.* Partial; \*\**p* < 0.01, Mann–Whitney *U* test). **(B)** Schematic illustration of the injury models and nerve-specific blockage used in the study. **(C)** Reversal of PCI-induced tactile hypersensitivity by intraplantar injection of capsaicin (10 μg) and QX-314 (6 mM in 20 uL of Saline). Data represent mean ± SEM (n=11-14 mice per group); \*\**p* < 0.01, Mann–Whitney *U* test. **(D)** Blockage of Aβ fibers (0.3 μg flagellin / 6 mM QX-314) did not alter PCI-induced tactile hypersensitivity. Data represent mean ± SEM (n=13-14 mice per group); Mann–Whitney *U* test. **(E)** 50% withdrawal threshold of ipsilateral paw following PCI in WT and TRPV1-KO mice. Both groups showed long-term tactile hypersensitivity. Data represent mean ± SEM (n=8-10 mice per group); Mann–Whitney *U* test. PCI, Partial sciatic nerve crush injury; BL, Baseline; WT, Wild type; KO, Knockout.

### 3.2 Nav1.8-positive nociceptor terminals partially remain after PCI

Following sciatic nerve PCI and FCI, we mapped denervation and reinnervation in the ipsilateral plantar hind paw using Thy1-YFP mice for Aβ-fiber and Nav1.8-tdTomato mice for Aδ/C-fibers [28; 58]. These mouse lines have been previously validated for distinguishing nociceptors and mechanoreceptors [31].

To visualize the anatomical distribution of nerve terminals, we performed whole-mount tissue clearing of the plantar hind paw skin (**Fig. 2A**). Thy1^+^ fibers showed complete denervation by day 7 and full reinnervation by day 30 in both crush models (**Figs. 2B-D**). In contrast, a significant proportion of Nav1.8^+^ fibers remained at day 7 post-PCI, whereas complete denervation was observed by day 7 after FCI (**Figs. 2E-G**). By day 30, reinnervation of Nav1.8^+^ fibers evident, but their density remained significantly lower after FCI than after PCI and the contralateral hind paw (**Figs. 2E-G**). These results indicate that the density of both spared and reinnervated nociceptor terminals is higher after PCI than after FCI.

**Figure 2.**
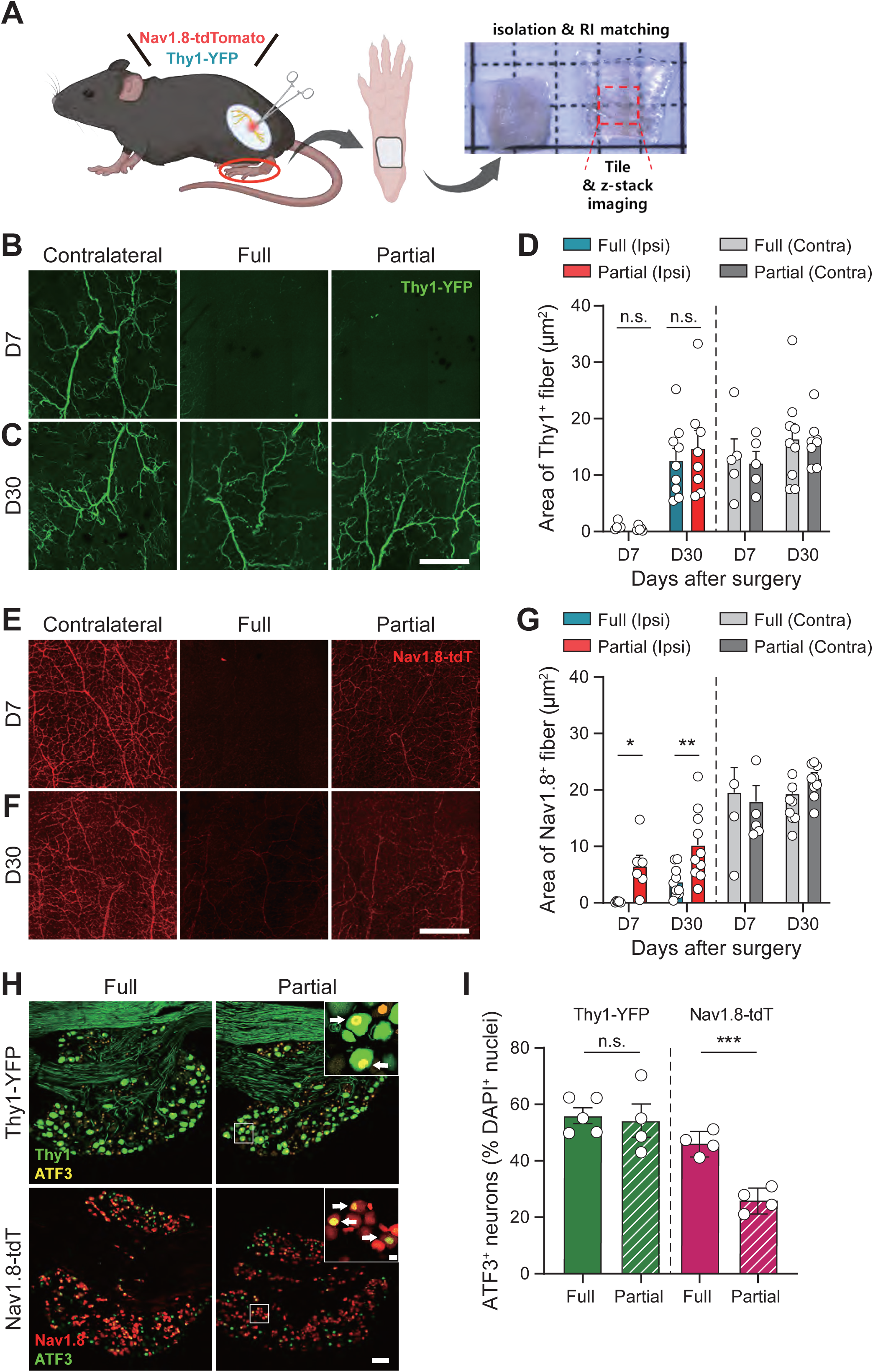
Nav1.8+ nociceptive nerve terminals remain following partial crush injury. **(A)** Schematic illustration of the whole mount imaging following both crush injuries in Thy1-YFP and Nav1.8-tdTomato mice. **(B-D)** Denervation and re-innervation pattern of Thy1^+^ fibers on day 7 and 30 following both crush injuries. Scale bar, 500 µm. Data represent mean ± SEM (D7 n=5 mice per group, Mann–Whitney *U* test; D30 n=8-9 mice per group, two-tailed unpaired *t*-test). **(E-G)** Denervation and re-innervation pattern of Nav1.8^+^ fibers at day 7 and 30 following both crush injuries. Scale bar, 500 µm. Data represent mean ± SEM (D7 n=5-6 mice per group, \**p* < 0.05, two-tailed unpaired *t*-test; D30 n=10 mice per group, \*\**p* < 0.01, two-tailed unpaired *t*-test). **(H, I)** Representative ATF3 images and average percentage of ATF3^+^ L4 DRG neuron on days 7 after both crush injuries in Thy1-YFP mice and Nav1.8-tdTomato mice. Arrow indicates ATF3^+^ neuron. Scale bar, 100 µm. Data represent mean ± SEM (n=4-5 mice per group, \*\*\**p* < 0.001, two-tailed unpaired *t*-test).

Next, to assess the extent of nerve injury, we quantified activating transcription factor 3 (ATF3) expression, a well-established marker of neuronal damage [74]. Consistent with the cutaneous innervation patterns in the hind paw (**Figs. 2A-G)**, the fraction of Thy1^+^ neurons expressing ATF3 did not differ between two crush injuries (**Figs. 2H, I**). In contrast, smaller fraction of Nav1.8^+^ neurons were ATF3^+^ after PCI than after FCI (**Figs. 2H, I**), suggesting greater sparing of Nav1.8^+^ nociceptors in PCI.

### 3.3 Miswiring of nociceptive fibers to mechanosensory target organ is not observed following both crush injuries

Recent studies link abnormal terminal connectivity to the development neuropathic pain [37; 79], with particular attention to the miswiring of nociceptors to mechanosensory end organs such as Merkel cells [29; 31; 40]. We therefore examined reinnervation of Nav1.8^+^ and Thy1^+^ nerve terminals into Merkel cells following crush injuries. The number of K8^+^ Merkel cells remained unchanged across injury types (**Supplemental Fig. 2**). Thy1^+^ fibers were fully denervated and subsequently reinnervated into the Merkel cells following both crush injuries (**Figs. 3A, B**), and Nav1.8^+^ fibers did not innervated Merkel cells at either time point, with no evidence of aberrant targeting (**Figs. 3C, D**). Quantitative analysis of Merkel cell innervation (**Supplemental Fig. 3**) revealed no significant changes in overlap volume or proximity between the Nav1.8^+^ fibers and Merkel cells either crush injuries (**Supplemental Figs. 3A-D**). In contrast, Thy1^+^ fibers consistently showed a greater overlap and shorter distances from the Merkel cell compared to Nav1.8^+^ fibers (**Supplemental Figs. 3E-H**). Together, these findings indicate that reinnervation occurs without misdirection, suggesting that miswiring is unlikely to contribute to PCI-induced hypersensitivity.

**Figure 3.**
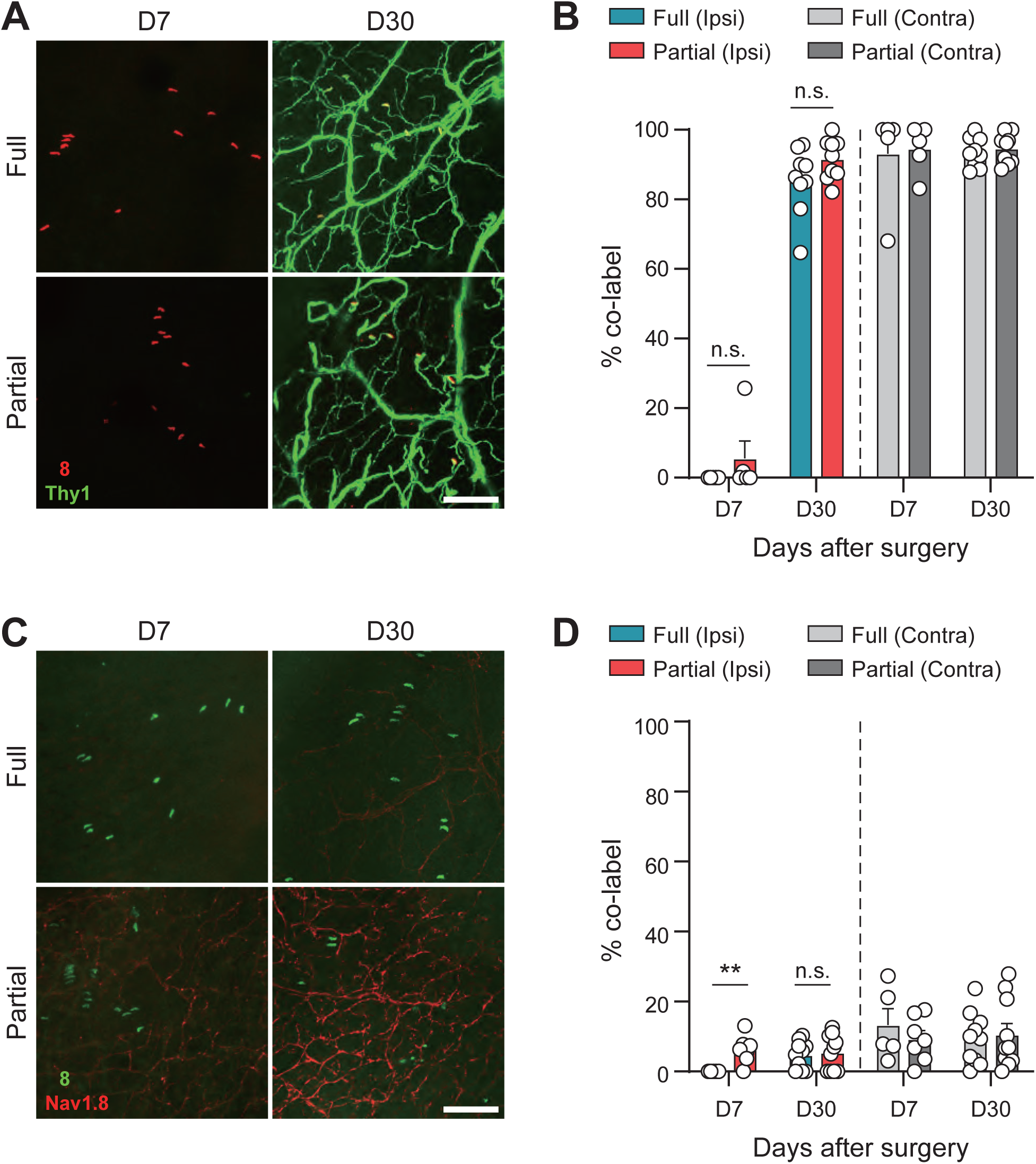
Thy1+ fibers fully innervate into Merkel cells following both crush injuries, whereas Nav1.8+ fiber do not. **(A)** Representative images of ipsilateral hind paw skin on days 7 and 30 after both crush injuries in Thy1-YFP mice. Merkel cells were stained with K8 (red). Scale bar, 100 µm. **(B)** Percentage of Thy1^+^ fibers co-labeled with the Merkel cells on day 7 and 30. Thy1^+^ fibers were fully denervated at day 7 and fully re-innervated at day 30 in both injury groups. Data represent mean ± SEM (D7 n=5-6 mice per group, Mann–Whitney *U* test; D30 n=9 mice per group, two-tailed unpaired *t*-test). **(C)** Representative images of ipsilateral hind paw skin on days 7 and 30 after both crush injuries in Nav1.8-tdTomato mice. Scale bar, 100 µm. **(D)** Percentage of Nav1.8^+^ fibers co-labeled with the Merkel cells on day 7 and 30. There was no significant increase in the mois-innervation of Nav1.8^+^ fibers to the Merkel cells in both injury groups. Data represent mean ± SEM (D7 n=5-7 mice per group, \*\**p* < 0.01, Mann–Whitney *U* test; D30 n=10 mice per group, Mann–Whitney *U* test).

### 3.4 Remaining cutaneous sensory neurons exhibit increased excitability after PCI

Sensory neuron hyperexcitability is a hallmark of neuropathic pain [33; 80; 88]. Given that Nav1.8^+^ neurons were more likely to remain after PCI (**Figs. 2-4**), we hypothesized that remaining nociceptor’s intrinsic excitability contributes to tactile hypersensitivity. To test this, we performed whole-cell patch clamp recordings from DRG neurons retrogradely labeled by injecting a fluorescent dye – DiI – into the ipsilateral hind paw on day 6 post-injury (**Fig. 4A**), allowing to distinguish remaining neurons from those that were injured. Recordings were conducted from small- (< 25 μm), medium- (25-35 μm), and large-diameter (> 35 μm) DRG neurons [30; 57].

**Figure 4.**
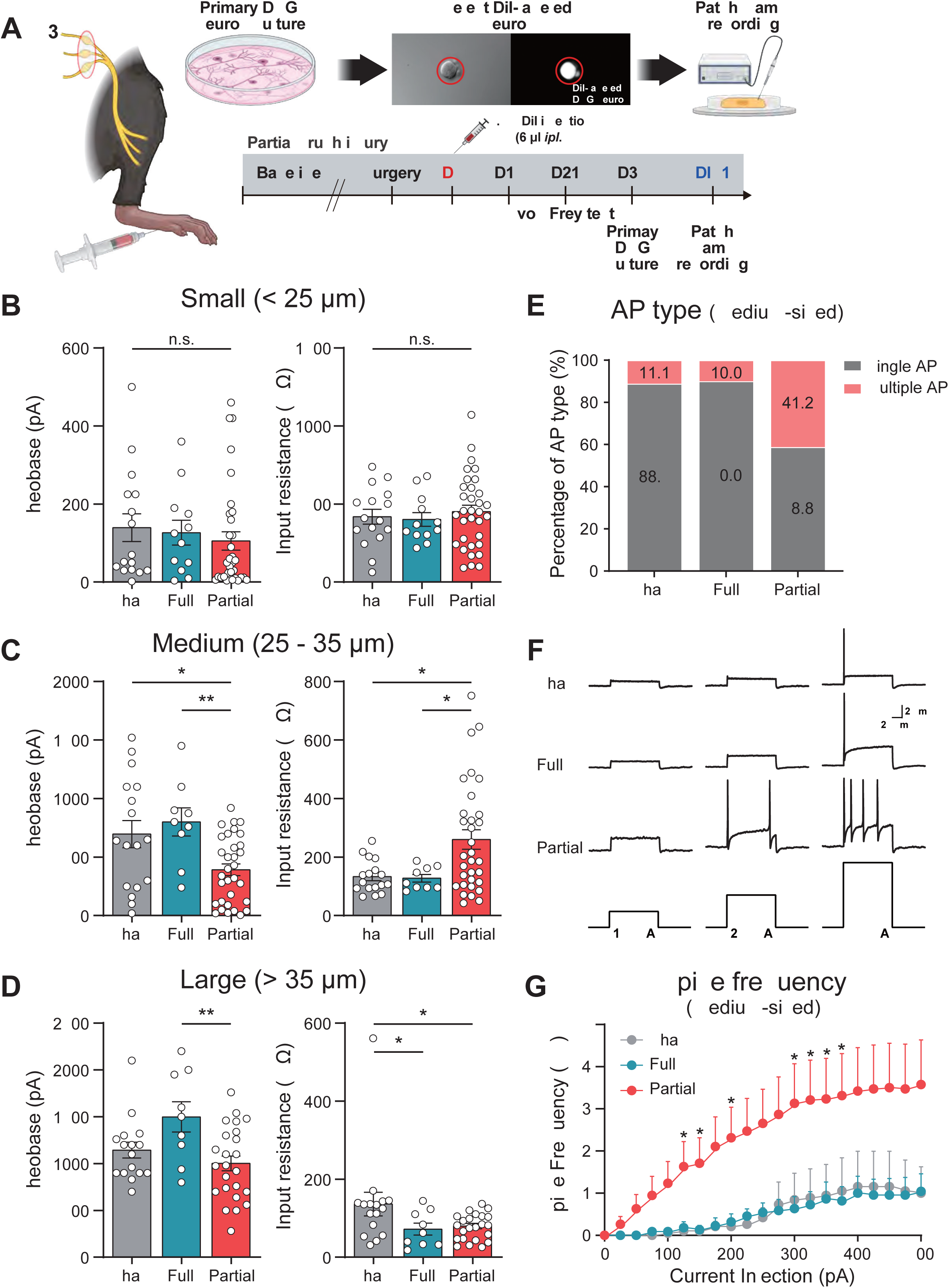
Remaining nociceptive DRG neurons are hyperexcitable after PCI. **(A)** Schematic illustration of DiI injection in mouse ipsilateral hind paw and timeline of behavioral assessment. **(B-D)** Rheobase and input resistance from small- (< 25 μm), medium- (25-35 μm) and large-diameter (> 35 μm) DiI^+^ DRG neurons 30 days after Sham, FCI, and PCI. Data represent mean ± SEM (small-diameter: rheobase n=12-33 DRG neurons per group, Mann–Whitney *U* test; input resistance n=12-33 DRG neurons per group, one-way ANOVA with Tukey multiple comparison) (medium-diameter: rheobase n=9-32 DRG neurons per group, \**p* < 0.05, \*\**p* < 0.01, Mann–Whitney *U* test; input resistance n=9-32 DRG neurons per group, \**p* < 0.05, Mann– Whitney *U* test) (large-diameter: rheobase n=9-24 DRG neurons per group, \*\**p* < 0.01, two-tailed unpaired *t*-test; input resistance n=9-24 DRG neurons per group, \**p* < 0.05, Mann–Whitney *U* test). **(E)** Percentage of AP type in medium-diameter DRG neurons after PCI. **(F)** Representative traces of APs following current injection. **(G)** Spike frequency in medium-diameter DRG neurons after Sham, FCI, and PCI. Data represent mean ± SEM. (n=19-38 DRG neurons per group, #, statistical significance compared to Sham; $, compared to FCI; *, compared to both Sham and FCI, Mann–Whitney *U* test)

To assess the intrinsic excitability of the DiI-labeled DRG neurons, we measured rheobase and input resistance. Medium-diameter neurons in PCI mice exhibited a significantly lower rheobase than Sham and FCI groups (**Figs. 4B-D**). Similarly, input resistance was significantly higher in the PCI group than in Sham or FCI groups (**Figs. 4B-D**), indicating increased excitability. Additionally, DRG neurons from PCI exhibited a higher proportion of multiple action potential (AP) firing than Sham or FCI (**Fig. 4E**) and showed elevated AP frequencies during graded current steps (0-500 pA) (**Figs. 4F, G**). Furthermore, single-cell RT-PCR confirmed expression of nociceptor markers (TRPV1, CGRP, and Nav1.8) in medium-diameter DRG neurons (**Supplemental Fig. 4**). These results indicate that spared medium-diameter cutaneous DRG neurons enriched for nociceptor markers exhibit intrinsic hyperexcitability after PCI, while consistent changes were not detected in small- or large-diameter neurons under our recording conditions.

### 3.5 Aδ- and C-fiber stimulation elicits enhanced dorsal horn responses after PCI

In both humans and mice, transcutaneous electrical stimulation at frequencies of 2000 Hz, 250 Hz, and 5 Hz preferentially activates Aβ-, Aδ-, and C-fibers, respectively [26; 45; 52; 53; 86]. To assess how spinal responses to fiber-biased stimulation, we quantified dorsal horn phosphorylated ERK (pERK) as a readout of neuronal activation [22; 32]. On day 30 after sham, FCI or PCI, stimulation was delivered to the ipsilateral hind paw under anesthesia (**Fig. 5A**). We confirmed that 2000 Hz stimulation primarily induced pERK expression in large-diameter DRG neurons, whereas 250 Hz and 5 Hz stimulations predominantly activated pERK in medium- to small-diameter DRG neurons (**Supplemental Fig. 5**).

**Figure 5.**
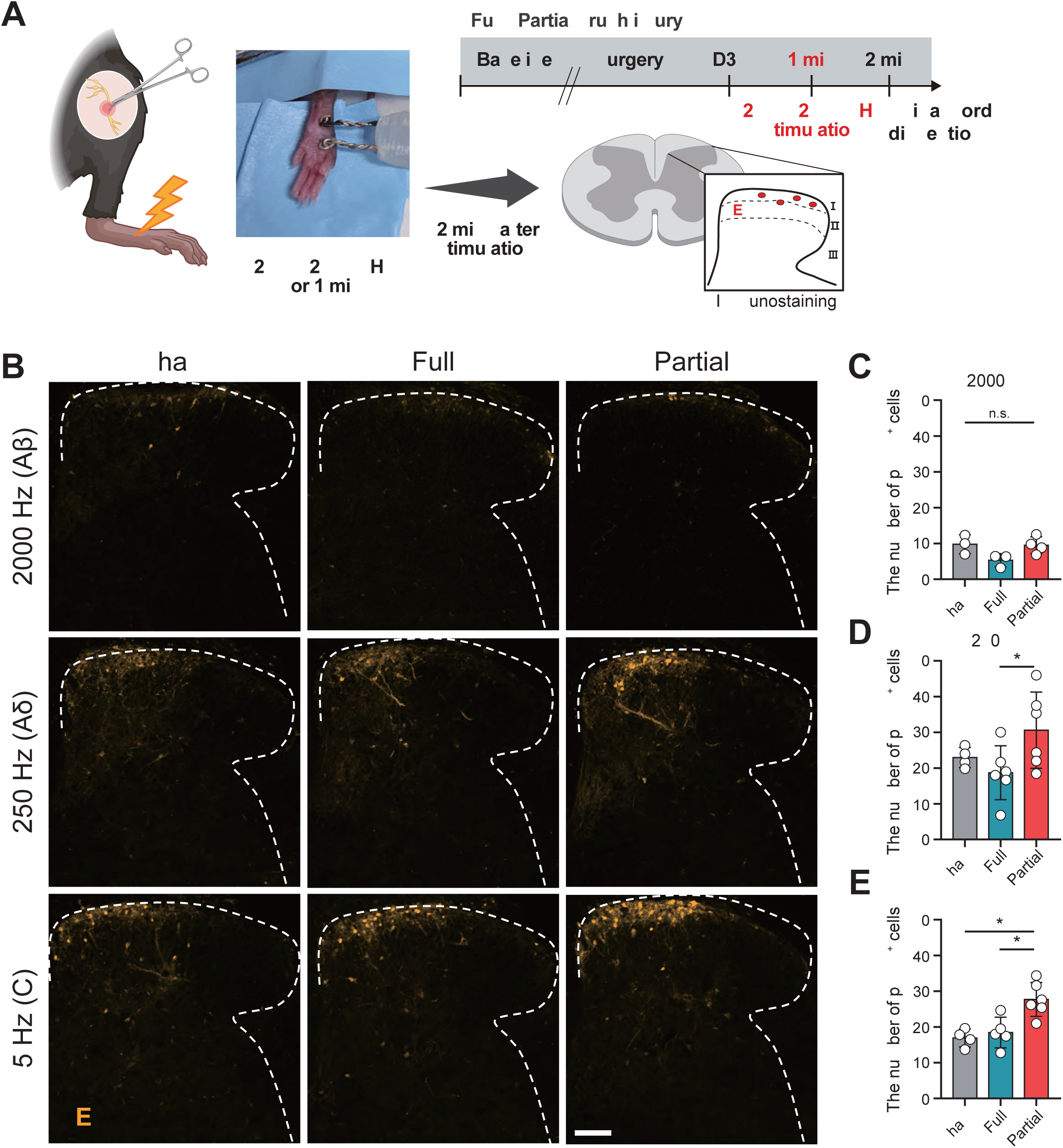
Aδ and C fiber electrical stimulation of the hind paw significantly increases pERK expression in the spinal dorsal horn. **(A)** Schematic illustration of the electrical stimulation for fiber type specific activation. **(B)** Representative images of pERK in the ipsilateral spinal dorsal horn after electrical stimulation of the ipsilateral hind paw with each of the three frequencies (2000, 250, and 5 Hz) on day 30 after surgery. **(C-E)** Quantitative analysis of pERK^+^ cells in the ipsilateral spinal dorsal horn. Scale bar, 100 µm. Data represent mean ± SEM (2000 Hz n=3-4 mice per group, Mann-Whitney *U* test; 250 Hz n=4-6 mice per group, \**p* < 0.05, one-way ANOVA with Tukey’s multiple comparison; 5 Hz n=4-6 mice per group, \**p* < 0.05, \*\**p* < 0.01, one-way ANOVA with Tukey’s multiple comparison)

In the dorsal horn, 2000 Hz stimulation did not measurably induce pERK in the spinal dorsal horn across all groups (**Figs. 5B, C**). In contrast, 250 Hz and 5 Hz stimulation led to a significant increase in pERK expression, particularly greater after PCI than after Sham or FCI (**Figs. 5B, D, E**). These findings indicate that dorsal horn input is potentiated by Aδ/C-, but not Aβ-, stimulation after PCI.

### 3.6 Early high-dose capsaicin-induced ablation of spared nociceptor terminals prevents the development of tactile hypersensitivity

Given evidence that nociceptor terminals persist after PCI and that spared cutaneous DRG neurons exhibit increased excitability, we hypothesized that early targeting of these neurons could prevent the development of long-term tactile hypersensitivity. To test this, we employed high-dose capsaicin subcutaneous injection, which is known to induce degeneration of TRPV1+ neuron terminal and alleviate neuropathic pain in humans and mice [3; 10; 17; 35].

A single high-dose of capsaicin (0.5 mg/mL, total 10 μg) or vehicle was injected into the ipsilateral hind paw on day 3 post-PCI (**Fig. 6A**). By day 7, Nav1.8^+^ terminals were significantly reduced in capsaicin-treated mice (**Figs. 6B, C**), while ATF3 expression in DRG neurons remained unchanged (**Supplemental Fig. 6**). Pinprick recovery was transiently delayed but returned to baseline by day 15 (**Fig. 6D**). Similarly, thermal hyposensitivity was observed 4 days after capsaicin injection and resolved by day 18 (**Fig. 6E**). Interestingly, tactile hypersensitivity failed to develop in capsaicin-treated mice, persisting for over >27 days post-capsaicin treatment (**Fig. 6F**). Notably, reinnervation of nociceptor terminals was comparable between vehicle- and capsaicin-treated groups at day 30 post-PCI (**Supplemental Fig. 7**), indicating that early ablation effectively prevents pain without impairing nerve subsequent reinnervation.

**Figure 6.**
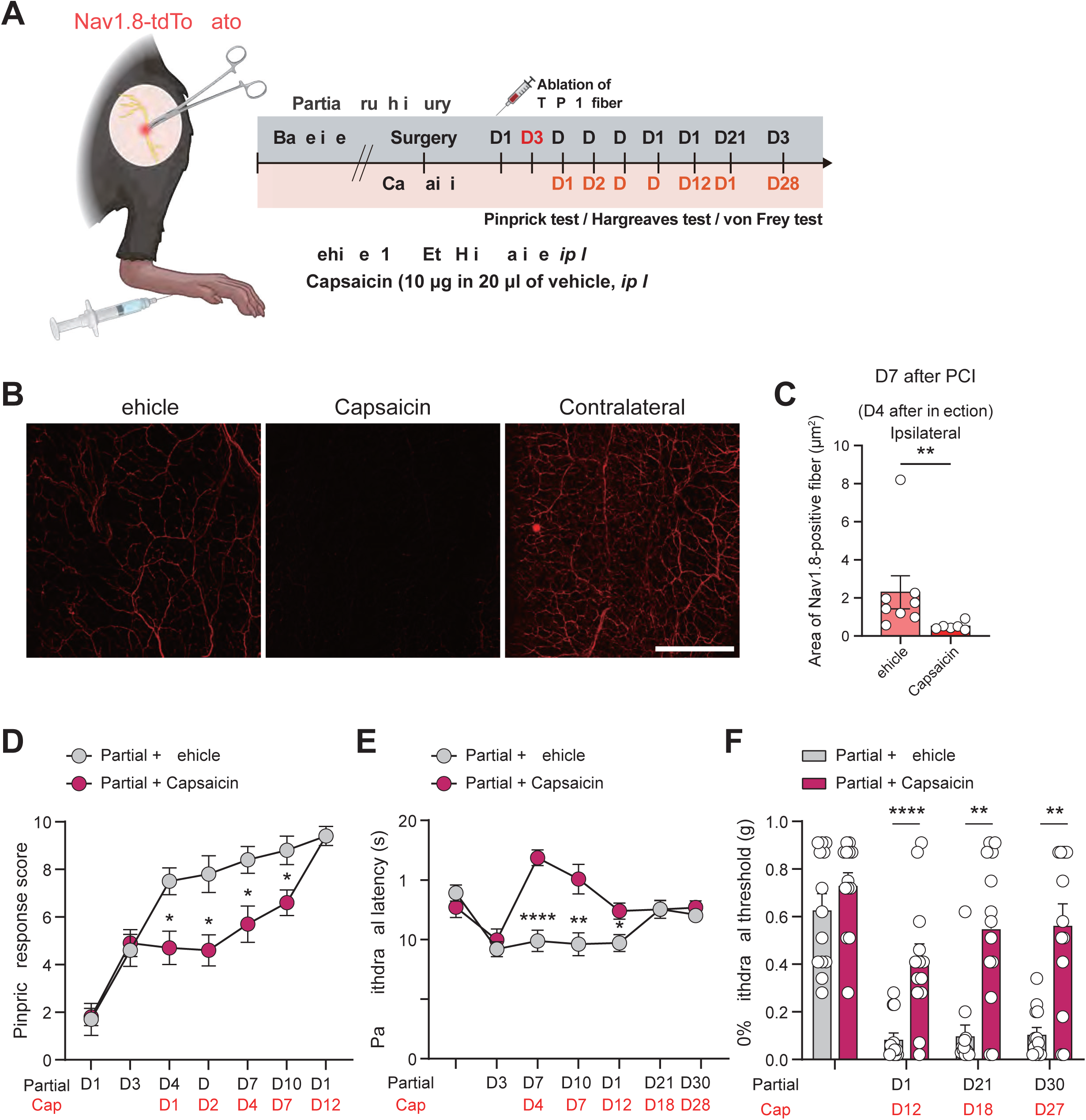
Chemical ablation of TRPV1+ fibers acutely after PCI relieves mechanical and thermal hypersensitivity. **(A)** Schematic illustration of the capsaicin-induced ablation of TRPV1^+^ fibers in PCI. **(B, C)** Denervation pattern of Nav1.8^+^ fibers 4 days after intraplantar injection of capsaicin or vehicle in PCI. n=6-8 per group. \*\**p* < 0.01, Mann–Whitney *U* test. Scale bar, 500 µm. **(D)** Pinprick response score (n=10 per group, \**p* < 0.05, Mann–Whitney *U* test), **(E)** withdrawal latency (n=9 per group, \**p* < 0.05, \*\**p* < 0.01, \*\*\**p* < 0.001, two-tailed unpaired *t*-test), **(F)** 50% withdrawal threshold (n=13-14 per group, \*\**p* < 0.01, \*\*\**p* < 0.001, Mann–Whitney *U* test) of ipsilateral hind paw after intraplantar injection of capsaicin (10 μg of capsaicin in 20 μL of vehicle) or vehicle (10% EtOH in saline) in PCI. **(G, H)** Re-innervation pattern of CGRP^+^ nociceptive fibers at day 30 following PCI in both groups. Scale bar, 500 µm. Data represent mean ± SEM (n=7-8 mice per group, two-tailed unpaired *t*-test). CGRP, calcitonin-related gene peptide.

Capsaicin administered on day 15, after tactile hypersensitivity had already developed, produced temporary relief, for 10 days post-injection (**Supplemental Fig. 8**). These results suggest that the sensitization of spared nociceptors during the degeneration period is critical for the development of long-term tactile hypersensitivity following PCI, and that early intervention against these afferents may help preventing the transition to chronic pain.

## 4. Discussion

Our findings reveal that spared nociceptive fibers are critical for the persistence of tactile hypersensitivity in sciatic nerve crush injury. Using Thy1-YFP and Nav1.8-tdTomato transgenic mice, we found that Nav1.8^+^ nociceptor terminals remained after PCI. Pharmacological silencing of TRPV1⁺ fibers with capsaicin/QX-314 significantly attenuated tactile hypersensitivity, confirming the functional contribution of nociceptors. Electrophysiological recordings showed hyperexcitability in medium-diameter sensory neurons, and increased pERK-positive cell counts in the spinal dorsal horn indicated increased central input. Lastly, local, high-dose capsaicin applied to the paw prevented the development of PCI-induced hypersensitivity when delivered early, underscoring the potential value of timing in targeting spared nociceptive fibers.

### 4.1 Transgenic labeling and pharmacological silencing reveal differential susceptibility of nociceptive and non-nociceptive fibers to nerve crush injury

The DRG contains a heterogeneous neuronal population, and conventional markers or behavioral assessment have limitations in distinguishing nociceptive from non-nociceptive subtypes [50; 66; 75]. We therefore employed two complementary anatomical and functional strategies: 1) we used Nav1.8-tdTomato mice to label nociceptors [58] and Thy1-YFP mice for Aβ mechanoreceptors [28], and 2) QX-314 co-applied with fiber-selective agonists to silence targeted subtypes.

Our data revealed a significant difference in the extent of neuronal damage between PCI and FCI by day 7 post-injury (**Fig. 2**). Thy1^+^ Aβ fibers were completely denervated in both models by this timepoint (**Figs. 2B-D**), as evidenced by comparable loss of terminal innervation and upregulation of ATF3 expression in DRG neurons (**Figs. 2H, I**). In contrast, Nav1.8^+^ nociceptive neurons were significantly less affected by PCI (**Figs. 2E-G**, **Figs. 2H, I**). These findings align with previous reports that large-diameter myelinated fibers are more susceptible to nerve compression injury than smaller fibers [21; 56; 69], likely reflecting the anatomical organization of large fibers into fewer, larger fascicles, which makes them more prone to injury [55; 73]. Functionally, intraplanar QX-314 with capsaicin revealed that Trpv1^+^ nociceptors are involved in tactile hypersensitivity following PCI (**Figs. 1C, D**). Prior studies indicated model-specific differences in fiber-type contributions to neuropathic pain; for instance, chronic constriction injury-induced pain is relieved by QX-314 with flagellin [85], while spared nerve injury-induced pain is alleviated by QX-314 with capsaicin [11]. Notably, our findings indicate that PCI-induced pain shares mechanistic similarities with SNI, as both appear to rely on the activity of spared TRPV1⁺ nociceptors.

Histological and pharmacological approaches cannot fully resolve Nav1.8^+^ versus TRPV1^+^ contributions. Nav1.8^+^ neurons encompass both peptidergic and non-peptidergic nociceptors and even some C-LTMRs or prurieceptors, whereas TRPV1 expression is mostly enriched in peptidergic subset [39; 60; 64]. Notably, our data demonstrate that high-dose capsaicin treatment significantly reduces the Nav1.8^+^ terminals (**Figs. 6B, C**), indicating substantial overlap. Dissecting the relative roles of these subtypes will require additional tools, but current data support a nociceptor-centric mechanism for PCI pain.

### 4.2 Denervation and regeneration of nociceptive fibers is not predictive of pain in PCI

Denervation of intraepidermal nerve fibers (IENFs) has been regarded as a gold standard of diagnosing small-fiber neuropathies (SFN) and its regeneration may correlate with reduced pain severity [24; 41; 61; 71]. However, some studies report conflicting results, showing that impaired IENF regeneration does not necessarily correlate with neuropathic pain [63; 89]. Interestingly, our findings revealed that the FCI model, which exhibits no pain-like behavior, had lower innervation density of nociceptive fibers compared to the PCI model (**Fig. 2**). Moreover, Nav1.8^+^ nociceptive fibers showed impaired regeneration, as they were not fully restored by day 30 in either crush injury compared to the contralateral side (**Figs. 2F, G**). These data indicate that decreased IENF density and their impaired regeneration do not correlate with pain behaviors after the crush injury. Although, the tissue clearing of hind paw skin in this study labels all cutaneous nerve fibers, not specifically IENF, our previous study showed spared IENF in PCI models [42]. Our results therefore suggest that the presence of spared Nav1.8^+^ fibers play a critical role in PCI-induced pain, implying that pain behaviors are less associated with the degree of IENF loss or the extent of nerve regeneration after injury. Additionally, given that axonal swelling in IENF fibers is a pathological feature in SFN patients [15; 38], further investigation into morphological changes of nociceptive fibers is warranted.

### 4.3 Lack of mis-targeted nociceptive fiber innervation in PCI-induced tactile hypersensitivity

Recent studies highlight that active regeneration processes play a pivotal role in maintaining neuropathic pain [19; 31; 40; 81; 84]. Following nerve injury, damaged axons regenerate toward their target organs. However, injured axons may form micro-neuromas, or mis-innervate into inappropriate target organs [68; 87]. Additionally, uninjured axons can sprout into the Merkel cells in denervated territories, potentially causing pain [29; 31]. Unlike spared nerve injury, involving chronic denervation of tibial and peroneal nerves, crush injury represents a single insult. It results in acute denervation followed by reinnervation across the entire hind paw skin. Although we examined Merkel cell innervation by Nav1.8^+^ fibers to assess potential mis-targeting after injury, the crush injury model complicates the clear observation of mis-innervation due to widespread denervation and subsequent reinnervation across the entire hind paw territory. Nevertheless, our results suggest that miswiring of nociceptive fibers into mechanosensory target organs did not occur, thereby ruling out miswiring as a major contributor to tactile hypersensitivity in PCI.

### 4.4 Spared nociceptor sensitization may drive tactile hypersensitivity

Neuronal hyperexcitability of peripheral sensory neurons has been observed in various neuropathic pain models and is closely associated with pain development [14; 16; 36; 90]. In preclinical nerve injury models such as L5 spinal nerve transection (L5 SpNT), uninjured L4 DRG neurons, separated from the injured L5 DRG, exhibit increased excitability contributing to pain [14; 34]. Our patch clamp recordings revealed that remaining medium-diameter neurons exhibited hyperexcitability following PCI (**Fig. 4**). A methodological limitation is that we relied on tracer labeling, whereas our anatomical quantification of spared terminals used the Nav1.8 reporter; thus, we cannot one-to-one link the hyperexcitable, tracer-identified units to the reporter-defined spared terminals. Nevertheless, post hoc confirmation of Nav1.8 and CGRP expression in these medium-diameter neurons (**Supplemental Fig. 4**) supports a nociceptor identity. Together, these data suggest that spared nociceptors maintain hyperexcitability, potentially playing a key role in PCI-induced tactile hypersensitivity. Notably, small-diameter DRG neurons did not show hyperexcitability following PCI, likely reflecting heterogeneity within this class; which includes various neuronal types (i.e. C-LTMR, pruriceptors, proprioceptors, and thermoreceptors) [7; 46; 66; 75] that possess distinct physiological and molecular characteristics and responses to nerve injury. Renthal et al., demonstrated subtype-specific transcriptional changes across injury models [66]; however, molecular drivers of nociceptor sensitization after crush injury remain unidentified, highlighting a need for further investigation.

Nociceptors transmit noxious stimuli to second-order neurons in the superficial dorsal horn of the spinal cord. Here, synaptic strength is enhanced under neuropathic pain conditions, leading to central sensitization [5; 82]. Our results showed significantly increased pERK expression within the spinal dorsal horn following electrical stimulation of Aδ, C-fibers in the PCI group (**Fig. 5**). In contrast, the partial sciatic nerve ligation model shows increased pERK expression in response to A-fiber but not C-fiber stimulation [53]. This indicates that input regulation by fiber type varies depending on the injury model. However, as we did not measure neuronal excitability or synaptic transmission in spinal neurons, further investigation is required.

### 4.5 Application of topical capsaicin by targeting spared nociceptive fibers alleviates neuropathic pain

The topical application of capsaicin has demonstrated therapeutic effects in various disease conditions [6; 27]. Since nociceptive fibers exclusively express the TRPV1 channel, capsaicin, a TRPV1 agonist, has been considered a potential treatment for desensitizing nociceptors in chronic pain [3; 13]. It is important to note that capsaicin can induce acute pain behaviors [59; 67], but it also resolves neuropathic pain by degenerating peptidergic afferents in nerve terminals [62; 65; 70; 76-78]. Consistent with this, topical application of capsaicin degenerated Nav1.8^+^ nerve fibers in the skin and provided an analgesic effect (**Figs. 6A-F**). Interestingly, genetic knockout of the TRPV1 did not reduce tactile hypersensitivity post-PCI, indicating TRPV1 itself may not be the primary pain driver (**Fig. 1F**). Supporting this, inhibiting TRPV1 in peripheral terminals did not alleviate pain in the ION-CCI model, while capsaicin-induced ablation of TRPV1^+^ fibers produced analgesic effects [77]. Moreover, the analgesic effect of capsaicin is transient and reversible in mice and humans [1; 2; 72; 77; 78]. Our data revealed that topical application of capsaicin at day 15, after pain-like behaviors had already developed, provided temporary relief of tactile hypersensitivity lasting 10 days post-treatment (**Supplemental Fig. 8**). In contrast, capsaicin treatment on day 3, during nerve degeneration, prevented pain development even after nerve regeneration (**Supplemental Fig. 7**). These findings suggest that the timing of the treatment significantly influences its effects, with early intervention potentially offering more sustained benefits in preventing pain.

In summary, our study highlights that the spared nociceptors after the PCI are a key driver of tactile hypersensitivity. This research offers valuable insights into their potential contribution of spared nociceptors to chronic pain following nerve crush injury, suggesting these fibers represent important targets for future investigation and therapeutic development for neuropathic pain.

## Acknowledgements

The authors thank Dr. Pilhan Kim at Korea Advanced Institute of Science and Technology (KAIST) for providing Thy1-YFP mice, Dr. Sunghoe Chang at Seoul National University for providing tissue clearing solution and Dr. Alexander Davies at Oxford University for providing valuable feedback on editing the manuscript. This research was supported by the National Research Foundation of Korea (NRF) grant funded by the Korean government (MSIT) RS-2021-NR059709, RS-2023-00264409, RS-2024-00441103 (to S.B. Oh), RS-2023-00240792 (to H.W. Kim), RS-2023-00272846 (to S.W. Shim). All authors gave their final approval and agreed to be accountable for all aspects of the work.

## Author contributions

Conceptualization: S.W. Shim, Y.K. Lee, K. Lee, H.W. Kim and S.B. Oh; Methodology and investigation: S.W. Shim, Y.K. Lee, D.H. Roh, H.W. Kim, K. Lee; Resources: S.B. Oh; Writing (original draft): S.W. Shim, Y.K. Lee; Writing (editing): S.W. Shim, Y.K. Lee, K. Lee, H.W. Kim and S.B. Oh; Project supervision: K. Lee, H.W. Kim and S.B. Oh; Correspondence: H.W. Kim (co-corresponding) and S.B. Oh (Primary).

## Data availability

The authors confirm that the data supporting the findings of this study are available within the article and its Supplemental materials. Additional study and data details are available from the corresponding author, [S.B. Oh], on special request.

