## Supplementary Information (PDF) for "Spared Nav1.8-Positive Nociceptors Drive Persistent Tactile Hypersensitivity After Sciatic Nerve Crush Injury in Mice"

### Supplemental Tables

**Supplemental Table 1** primer sequence for single cell RT-PCR

| Target gene | scRT-PCR | | GenBank No. |
|---|---|---|---|
| | Outer primers (5' to 3') | Inner primers (5' to 3') | |
| GAPDH<br>(282 bp) | | (F) CCAGAACATCATCCCTGCAT<br>(R) GCATCGAAGGTGGAAGAGTG | NM_001289726.1 |
| TRPV1<br>Outer : 293bp<br>Inner : 115 bp | (F) CATCTTCACCACGGCTGCTTA<br>(R) CATCTTCACCACGGCTGCTTA | (F) GCGACCATCCCTCAAGAGTT<br>(R) ATACTCCTTGCGATGGCTGA | NM_001001445.2 |
| CGRP<br>Outer : 242 bp<br>Inner : 156 bp | (F) GCCTTTGAGGTCAATCTTGG<br>(R) CCTTCACCACACCTCCTGAT | (F) CACTCTCAGTGAAGAAGAAGTTCG<br>(R) GTCACACAGGTGGCAGTGTT | NM_001033954.3 |
| $Na_V1.8$<br>Outer : 311bp<br>Inner : 190 bp | (F) GCCACTTCTTCTGGGTTAACG<br>(R) GTACTTCTTCTGCTCCTCTGTCAT | (F) CGTTGCTATGGGCTACCTCG<br>(R) CCCGACAAAGAGATTCAGCG | NM_001205321.1 |

**Supplemental Table 2** Antibodies for immunohistochemistry

| Antibodies | Conjugate | Host | Company | Cat No. |
|---|---|---|---|---|
| TROMA-I (K8) | none | Rat | DSHB | AB_531826 |
| CGRP | none | Goat | Abcam | ab36001 |
| ATF3 | none | Rabbit | Novus Biological | NBP1-85816 |
| Phospho-p44/42 MAPK (Erk1/2) | none | Rabbit | Cell Signaling | 9101 |
| Rat IgG | Alexa fluor 488 | Donkey | Jackson Laboratory | 712-545-153 |
| Rat IgG | Cy3 | Donkey | Jackson Laboratory | 712-165-153 |
| Goat IgG | Alexa fluor 488 | Donkey | Jackson Laboratory | 705-545-003 |
| Rabbit IgG | Alexa fluor 488 | Donkey | Jackson Laboratory | 711-545-152 |
| Rabbit IgG | Cy3 | Donkey | Jackson Laboratory | 711-165-152 |

### Supplemental Figures

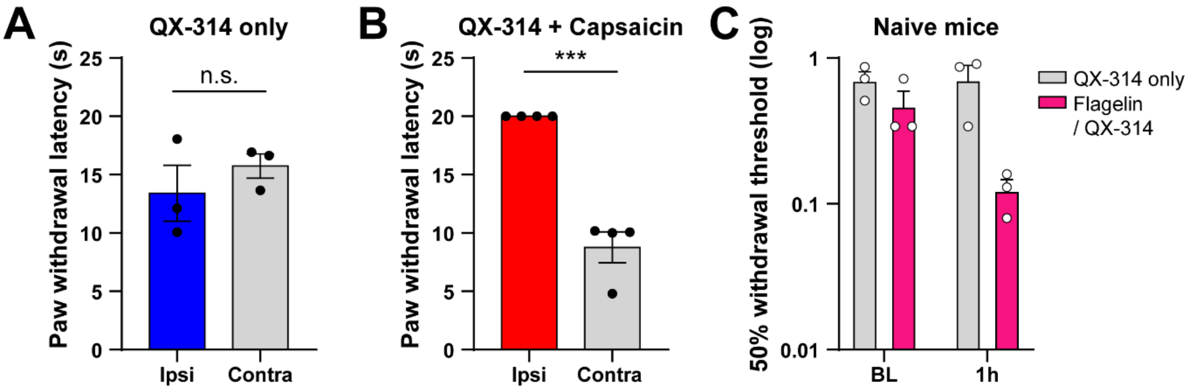


**Supplemental Figure 1. Co-application of QX-314 with capsaicin or flagellin produces fiber-selective blockade in naïve mice.**

**(A)** Paw withdrawal latency during thermal stimulation following intraplantar injection of QX-314 only (6 mM) N=3 mice per group. two-tailed unpaired *t*-test. **(B)** Paw withdrawal latency during thermal stimulation following intraplantar injection of QX-314 (6 mM) with capsaicin (10 µg). Application of QX-314 with capsaicin showed thermal insensitivity in naive mice. N=4 mice per group. $^{**}p < 0.01$, two-tailed unpaired *t*-test. **(C)** 50% withdrawal threshold of ipsilateral hind paw following intraplantar injection of QX-314 (6 mM) with flagellin (0.3 µg) in naïve mice. A fiber blockage in WT mice showed tactile hypersensitivity an hour after the application. N=3 mice per group. Data represent mean ± SEM.

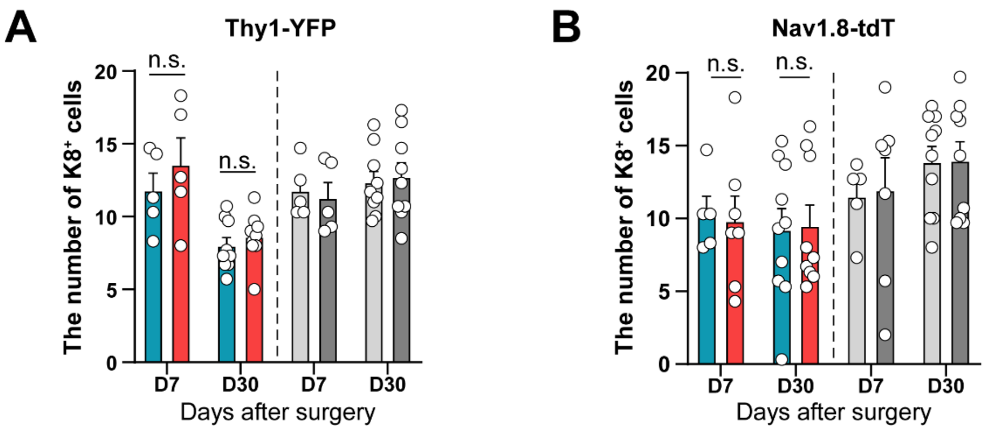


**Supplemental Figure 2. K8$^+$ Merkel cell counts are unchanged after PCI and FCI in Thy1-YFP and Nav1.8-tdTomato mice**

**(A)** The number of K8$^+$ Merkel cells on day 7 and 30 in Thy1-YFP mice. **(B)** The number of K8$^+$ Merkel cells on day 7 and 30 in Nav1.8-tdTomato mice. Data are presented as means ± SEM. two-tailed unpaired *t*-test.

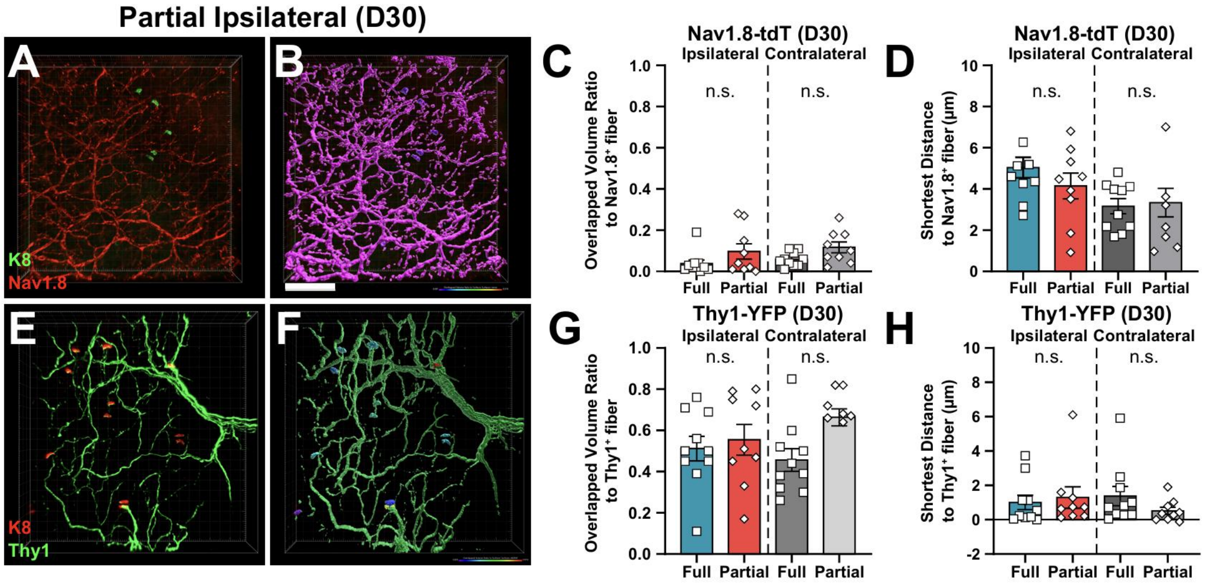


**Supplemental Figure 3. No evidence of nociceptor mistargeting to mechanosensory end organs following PCI.**

**(A)** Representative images of ipsilateral hind paw skin on day 30 after partial crush injury in Nav1.8-tdTomato mice. Merkel cells were stained with K8 (Green). **(B)** Representative 3D rendering image of Nav1.8$^{+}$ fibers (Red) and Merkel cell (Green) using IMARIS. **(C)** Overlapped volume ratio to Nav1.8$^{+}$ fibers from the Merkel cells. Data are presented as means ± SEM. Mann–Whitney *U* test. **(D)** Shortest distance to Nav1.8$^{+}$ fibers from the Merkel cells. Data are presented as means ± SEM. two-tailed unpaired *t*-test. **(E)** Representative images of ipsilateral hind paw skin on day 30 after partial crush injury in Thy1-YFP mice. Merkel cells were stained with K8 (Red). **(F)** Representative 3D rendering image of Thy1$^{+}$ fibers (Green) and Merkel cell (Red) using IMARIS. **(G)** Overlapped volume ratio to Thy1$^{+}$ fibers from the Merkel cells. Data are presented as means ± SEM. two-tailed unpaired *t*-test. **(H)** Shortest distance to Thy1$^{+}$ fibers from the Merkel cells. Data are presented as means ± SEM. Mann–Whitney *U* test.

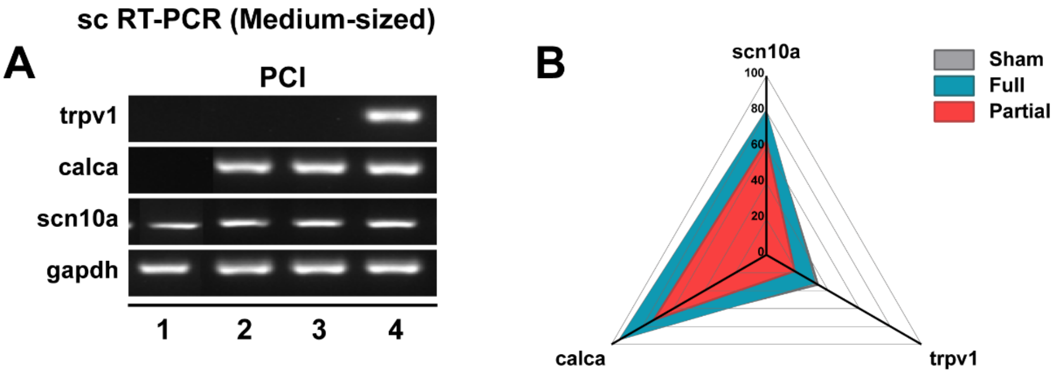


**Supplemental Figure 4. Single cell RT-PCR confirms expression of nociceptor markers in medium-diameter DiI-labeled neurons.**

**(A, B)** Single cell RT-PCR analysis for trpv1, calca and scn10a mRNA expression in medium-diameter DRG neurons after PCI.

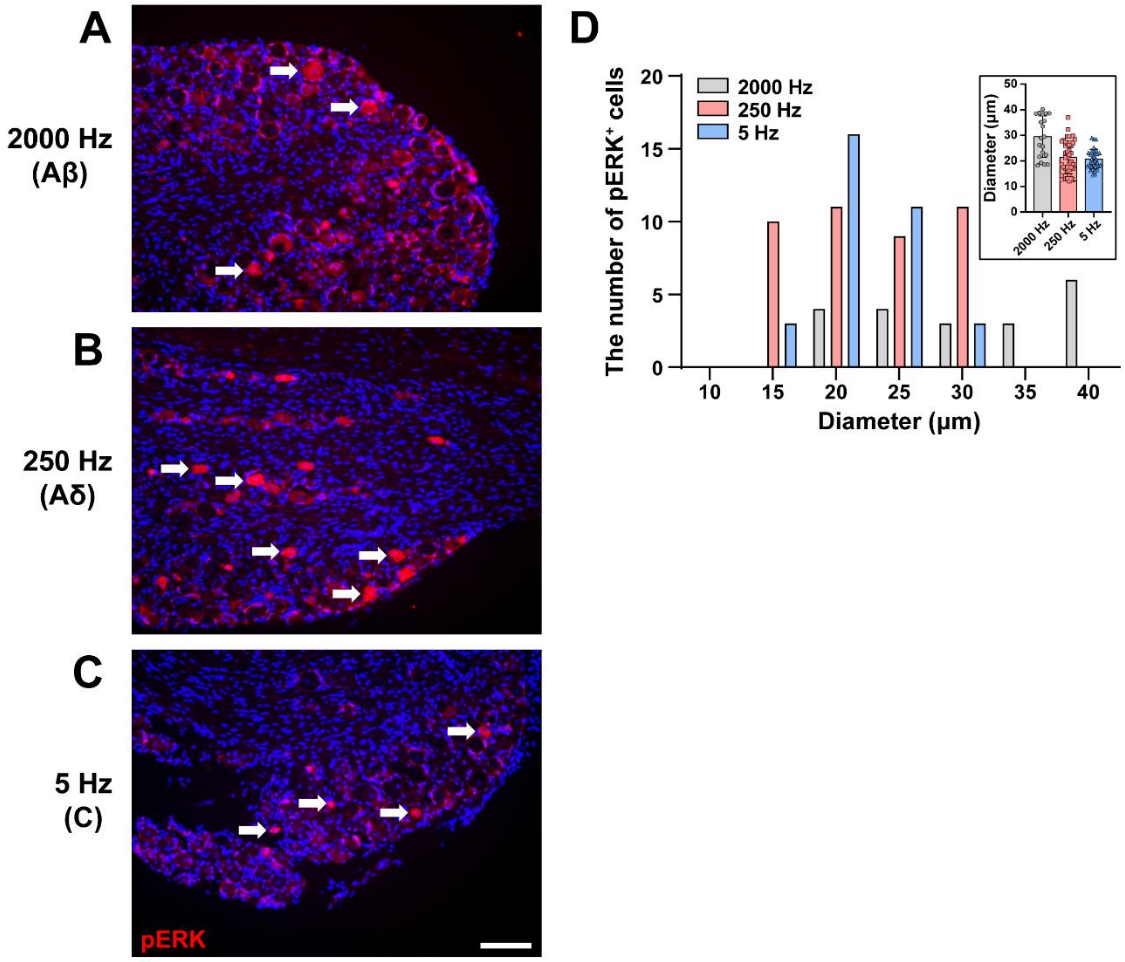


**Supplemental Figure 5. pERK expression patterns in DRG neuron after 2000, 250, and 5Hz electrical stimulation.**

**(A-C)** Representative images of pERK in the L4 DRG of Sham mice after electrical stimulation to the right hind paw with each of the three frequencies (2000, 250, and 5 Hz) Arrow indicates pERK-positive DRG neuron. **(D)** The cell size distribution of pERK$^{+}$ DRG neuron after electrical stimulation. Scale bar, 75 μm.

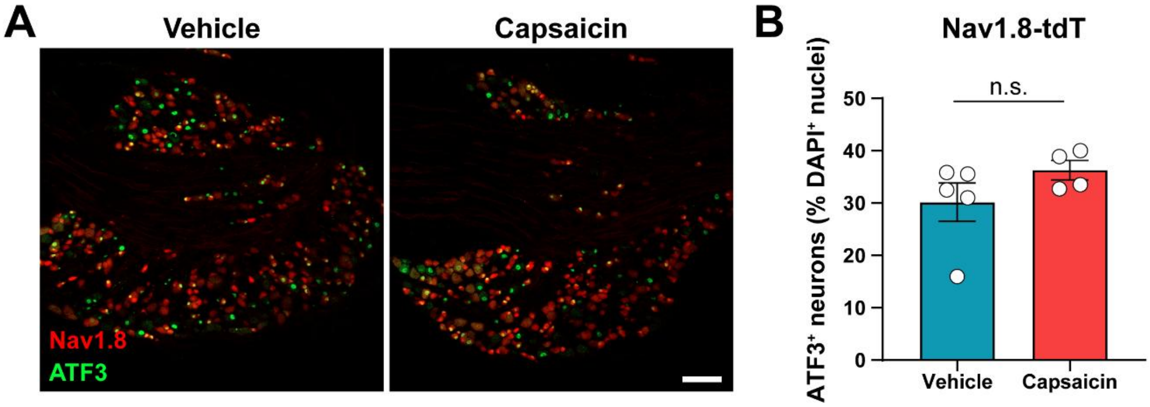


**Supplemental Figure 6. High-dose capsaicin injection does not alter ATF3 expression in DRG neurons**

**(A)** Representative ATF3 images of L4 DRG neuron **(B)** Average percentage of ATF3 expression in Nav1.8$^{+}$ neurons after intraplantar injection of Vehicle (10% EtOH in saline) or Capsaicin (10 µg of capsaicin in 20 µL of Vehicle) in PCI. Scale bar, 100 µm. Data represent mean ± SEM (n=4-5 mice per group, Mann–Whitney *U* test).

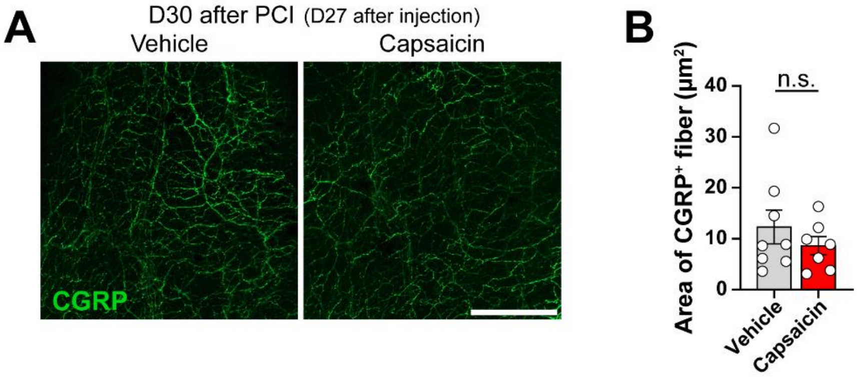


**Supplemental Figure 7. Reinnervation of CGRP$^+$ nociceptor terminals at day 30 after PCI is comparable between vehicle- and capsaicin-treated groups.**

**(A)** Representative images of CGRP$^+$ fibers in the ipsilateral hind paw 30 days after PCI (27 days after capsaicin injection.). **(B)** Quantitative analysis of CGRP$^+$ fibers in ipsilateral hind paw. Scale bar, 500 μm. Data represent mean ± SEM (n=7-8 mice per group). CGRP, Calcitonin-related gene peptide.

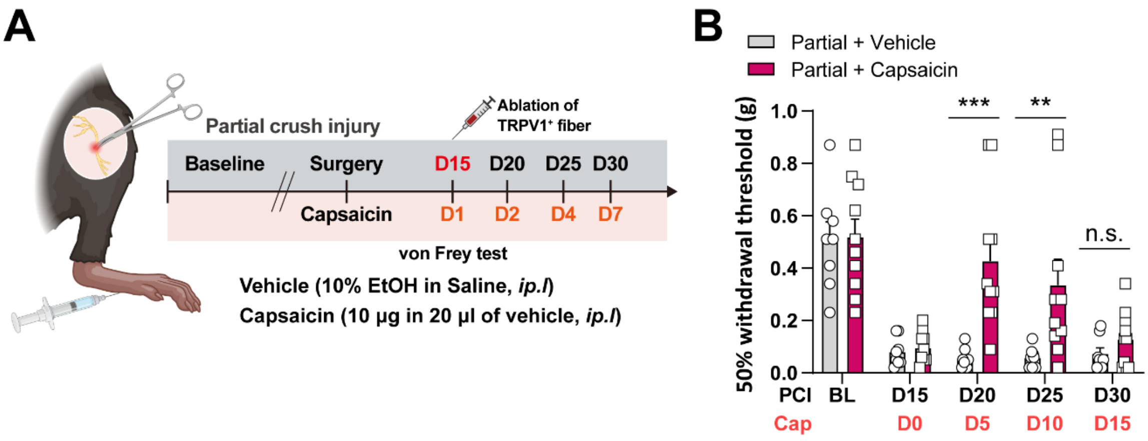


**Supplemental Figure 8. Late (day 15) ablation of TRPV1$^+$ terminals yields transient reversal of PCI-induced tactile hypersensitivity.**

**(A)** Schematic illustration of the capsaicin-induced ablation of TRPV1$^+$ fibers in PCI. **(B)** 50% withdrawal threshold of ipsilateral paw after intraplantar injection of capsaicin (10 µg of Capsaicin in 20 µL of Vehicle) or vehicle (10% EtOH in Saline) in PCI. N= 8-10 mice per group. Data are presented as means ± SEM. ***$p < 0.01$, ****$p < 0.001$. Mann–Whitney *U* test.
